# A low dimensional manifold of human exploratory behavior reveals opposing roles for apathy and anxiety

**DOI:** 10.1101/2023.06.19.545645

**Authors:** Xinyuan Yan, R. Becket Ebitz, Nicola Grissom, David P. Darrow, Alexander B. Herman

**Author notes:** Corresponding author: Alexander B. Herman, **Email:**. Denotes equal contribution (David P. Darrow, Alexander B. Herman). **Author Contributions:** Conceptualization, A.B.H, R.B.E, N.G, D.P.D; Methodology, X.Y, R.B.E, A.B.H; Formal Analysis, X.Y; Writing – Original Draft, X.Y, D.P.D, A.B.H; Funding Acquisition, A.B.H and N.G; Supervision, A.B.H, D.P.D. **Competing Interest Statement:** The authors declare no competing interests.

## Abstract

Exploration-exploitation decision-making is a feature of daily life that is altered in a number of neuropsychiatric conditions. Humans display a range of exploration and exploitation behaviors, which can be affected by apathy and anxiety. It remains unknown how factors underlying decision-making generate the spectrum of observed exploration-exploitation behavior and how they relate to states of anxiety and apathy. Here, we report a latent structure underlying sequential exploration and exploitation decisions that explains variation in anxiety and apathy. 1001 participants in a gender-balanced sample completed a three-armed restless bandit task along with psychiatric symptom surveys. Using dimensionality reduction methods, we found that decision sequences reduced to a low-dimensional manifold. The axes of this manifold explained individual differences in the *balance* between states of exploration and exploitation and the *stability* of those states, as determined by a statistical mechanics model of decision-making. Position along the balance axis was correlated with opposing symptoms of behavioral apathy and anxiety, while position along the stability axis correlated with the level of emotional apathy. This result resolves a paradox over how these symptoms can be correlated in samples but have opposite effects on behavior. Furthermore, this work provides a basis for using behavioral manifolds to reveal relationships between behavioral dynamics and affective states, with important implications for behavioral measurement approaches to neuropsychiatric conditions.

## Introduction

We often face the dilemma of whether to exploit a known option or explore potentially better alternatives. When reward contingencies are fixed, simple exploration-exploitation strategies such as repeating a rewarded choice or switching after a loss can lead to success. However, in dynamic, changing environments, more reflective of the real world, a diverse range of strategies are possible (1, 2). As the number of sequential decisions and outcomes in a dynamic environment increases, the potential complexity of the mix of these different strategies increases exponentially.

Affective states, including levels of motivation and anxiety (3–5) influence the mix of exploration and exploitation strategies, possibly through cognitive biases such as reward and uncertainty sensitivity (6, 7). For instance, anxious individuals may be driven to explore unknown options rather than exploit a rewarding one, while apathetic individuals may choose to stay with a decision even when it’s no longer rewarding. Changes in the balance of exploration-exploitation strategies have also been observed in several clinical neuropsychiatric conditions (4, 8), such as schizophrenia (9), addiction (10), and anxiety disorders (3).

To untangle the factors generating diverse exploration-exploitation behaviors and how they contribute to psychiatric symptoms, we recruited 1001 participants to complete a three-armed restless bandit exploration-exploitation task (Figure 1A) and psychiatric measurements of mood, motivation, and anxiety. We employed advances in dimensionality reduction methods to test whether human exploration-exploitation decision-making occupies a low-dimensional shape and whether such a shape can explain individual differences in psychiatric measures important for decision-making. Dimensionality reduction on personality traits has demonstrated utility in identifying the latent structures of psychological states (11–13) as well as neural states (14–16), but has not been applied to behavioral processes.

**Figure 1.**
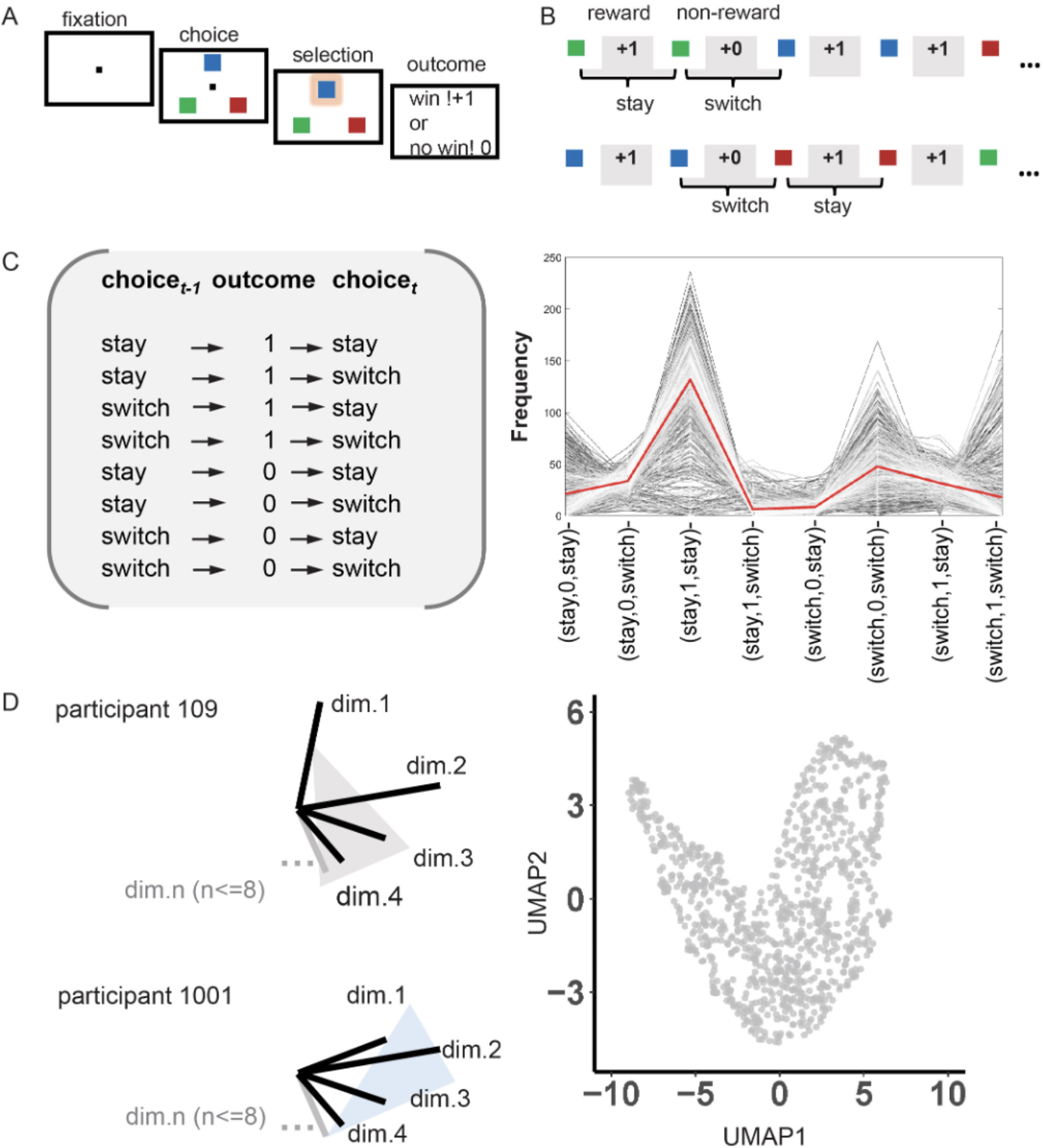
Dimensionality reduction of behavior on a three-armed restless bandit exploration-exploitation task. (A) three-armed restless bandit task. Participants chose one option from among the three targets to receive reward or non-reward feedback. Each target is associated with a hidden reward probability that randomly and independently changed throughout the task. (B) example choice and reward sequence and the definition of *stay* and *switch*. Specifically, *stay* was defined as choosing the same target as in the previous trial, while *switch* was defined as choosing a different target. “+1” denotes reward feedback, “+0” denotes non-reward feedback. (C) all possible choice-reward sequences (left) and the distribution (right) of individual differences in frequency for all decision patterns. The red line represents the mean value for each decision pattern across participants. (D) schematic high dimensional space of participants’ decision-making pattern (left) and the manifold-projected behavior (right).

Our results show that exploration and exploitation behaviors collapse to a low-dimensional manifold, which forms along dimensions corresponding to the balance and stability of decision-making states. The shape of this manifold explains how apathy and anxiety, which have been studied separately but rarely together, oppose each other in exploratory behavior.

## Results

Gender-matched adults (n=1001, age: range 18-54, mean ± SD = 28.446 ± 10.354 years; gender: 493 female) performed a restless 3-armed bandit (Figure 1A) (17, 18), and completed symptom surveys on mood, anxiety, and motivation (See methods and SI Section1, Table S1). In the bandit task, participants were presented with images of three playing cards and moved their cursor over their choice. The probability of reward associated with each card deck drifted randomly over time, encouraging participants to balance periods of exploitation, when they’ve found a good option, with periods of exploration, when they search for a better one.

### Dimensionality reduction of behavior on a three-armed restless bandit exploration-exploitation task

To capture the full richness of behavior in the bandit task, we formatted each participants’ trial-by-trial task data into sequences of choices to stay (repeat the choice on the last trial) or switch (choose a different option) and reward outcome for two consecutive trials ({choice_t-1_, outcome_t-1_, choice_t_}, Figure 1B and 1C). The behavioral data for each participant was then transformed into counts for each of these eight unique sequences. To determine whether behavior on this high-dimensional task can be reduced to a low-dimensional representation, we applied Uniform Manifold Approximation and Projection (UMAP) (19), a computationally efficient algorithm that can preserve both the local and global distances between data points in high-dimensional space, to learn the two dimensional manifold underlying the eight-dimensional behavioral data, see Methods for more algorithm details). Including additional reward-history did not affect the dimensionality reduction results (SI Section 2, Figure S1).

### A low dimensional manifold of individual differences in exploration-exploitation decision-making

We next asked how the manifold explains the underlying variation in exploration-exploitation decision-making in our sample. To reveal latent states of exploration and exploitation, and the dynamics of transitions between those states, we fit a Hidden Markov Model (HMM) to the sequences of decisions (18, 20). We mapped model-free (Figure 2A) and model-based (Figure 2B and 2C) behavioral measures onto the UMAP (all model-free and model-based measures, see SI Section 3, Table S2). We found that patterns of exploration and exploitation were indeed well-described by the UMAP projection, yielding clearly interpretable results. We then conducted correlation analyses between all indices and the axes of the manifold (i.e., UMAP1, the horizontal axis, and UMAP2, the vertical axis) to quantify the relationships.

**Figure 2.**
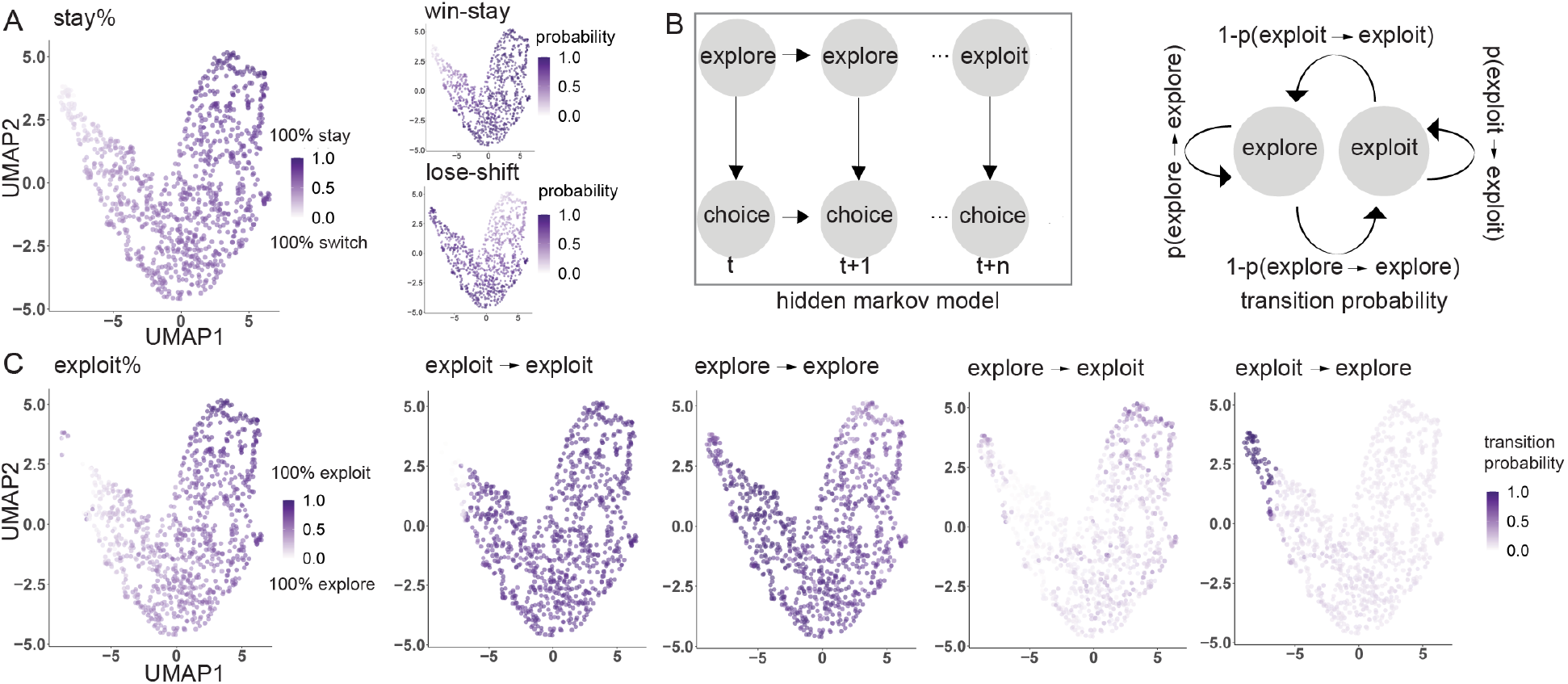
A low dimensional manifold of individual differences in exploration-exploitation decision-making. (A) individual differences in model-free measures of sequential behavior projected onto the manifold (percentage of stay, win-stay, and lose-shift decisions). (B) schematic of Hidden Markov Model (HMM) (left) and transition probabilities within and between states (right). (C) individual differences in HMM model-based measures of decision states and transition probabilities projected onto the manifold (from left to right, percentage of exploit states; probability of transition from: exploit t0 exploit, explore to explore, explore to exploit, and exploit to explore).

The horizontal-axis dimension of the manifold explained most of the variance in both model-free stay-switch and model-based exploration-exploitation behavior. Moving from left to right along the horizontal axis (UMAP1) of the manifold, the probability of switching decreased, as did the probability of exploration, while the probability of staying and the probability of exploiting increased, with a high correlation coefficient (Figure 2A and 2C, SI Section 4, Table S3 & Table S4, all p < 0.001, corrected by FDR p < 0.05). When we decomposed the behavior into the degree with which individuals stay after a reward (“win-stay’) and switch after reward omission (“lose-shift”), we found that the horizontal axis largely explained the win-stay behavior in the positive direction and the lose-shift behavior in the opposite direction.

### The energy landscape explaining manifold dimensions

To understand how the dynamics of exploration-exploitation decision-making map onto the manifold, we examined the probabilities of the different state transitions. We found that the axes of the manifold explained different features of the exploration-exploitation transition dynamics. The horizontal axis (UMAP1) was aligned with the overall probability of exploitation. It was positively associated with the probability of transitioning from explore to exploit, positively associated with exploit to exploit, negatively associated with exploit to explore, and negatively associated with explore to explore (all p < 0.001, all FDR-corrected p < 0.05; see Table S4). In contrast, the vertical axis (UMAP2) was negatively associated with the probability of remaining in either an exploit or an explore state, and positively associated with transitioning between states (Figure 3A, all p < 0.001, all FDR-corrected p < 0.05; see Table S4). We further analyzed the dynamics with the tools of thermodynamics (20). The stationary probability of occupying a given state is proportional to the depth of that state’s potential energy well. The probability of transitioning between states, in turn, is related to the height of the activation energy barrier separating them. A higher transition probability indicates that less activation energy is needed to cross the barrier, leading to a more unstable system. To further confirm our explanations for UMAP1 and UMAP2, we did correlation analyses between the UMAP score and the difference of the stationary distribution probabilities (i.e., the difference in energy that is associated with exploration and exploitation), as well as the activation energy, which captures the system’s stability. Results showed that UMAP1 more strongly tracked the difference of potential energy (r = 0.812, p < 0.001) for each state compared to UMAP2 (z-score = 21.971, p < 0.001). While UMAP2 more strongly tracked the activation energy (r = -0.228, p < 0.001) compared to UMAP1 (z-score = 3.125, p = 0.001) (Figure 3B). Thus, the relative depths of the energy wells corresponding to explore and exploit states was most closely associated with the horizontal axis (UMAP1), which we call the balance axis, whereas the height of the activation energy barrier between them best characterized the vertical axis (UMAP2), which we refer to as the stability axis (Figure 3C).

**Figure 3.**
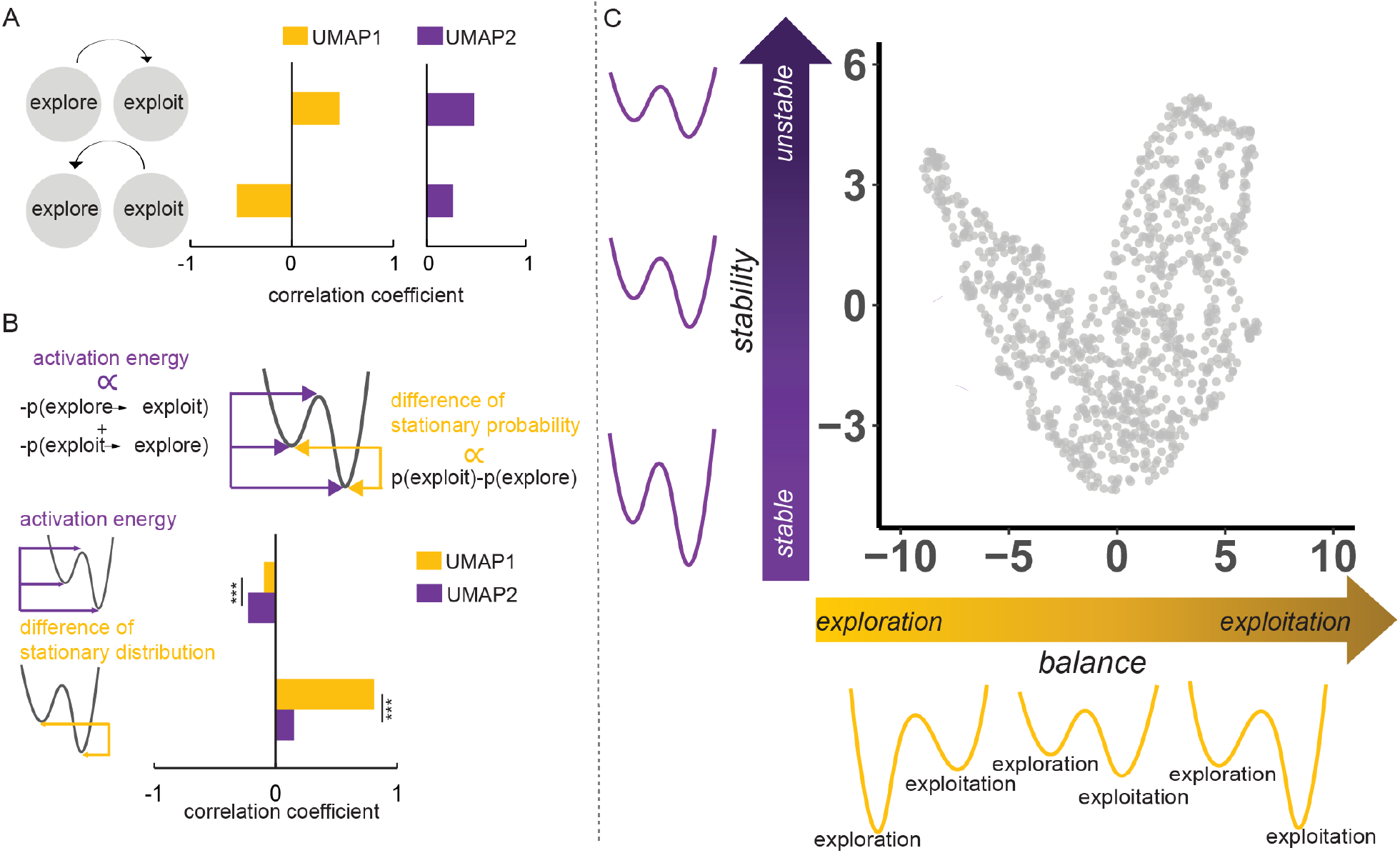
The energy landscape explaining manifold dimensions. (A) correlation coefficients between the state transitions and the manifold horizontal axis (UMAP1, yellow) and the vertical axis (UMAP2, purple), demonstrating that UMAP1 most closely tracked the exploration-exploitation balance, while UMAP2 was associated with state stability. (B) schematic (top) and correlation coefficients (bottom) showing that the UMAP1-balance association is explained by the difference in energy well depths (stationary distribution probabilities) of each state, while the UMAP2-stability association relates to the activation energy of state transitions. (C) schematic with manifold-projected data summarizing how the manifold is shaped by a horizontal dimension of exploration-exploitation state balance and a vertical dimension of state stability.

### Manifold projection of individual differences in anxiety and apathy subdomains

We next examined whether the latent manifold explained features of affect important for decision making. We conducted Spearman correlation analyses to reveal associations between the UMAP scores and psychiatric symptoms.

Our results showed that both the horizontal axis (UMAP1) of the manifold (rho = 0.117, p < 0.001) and vertical axis (UMAP2) (rho = 0.101, p = 0.001) correlated positively with apathy, as well as social apathy (subscale to measure the motivation deficits in social interaction) (horizontal axis, rho = 0.117, p < 0.001; vertical axis, rho = 0.058, p = 0.065). The behavioral apathy (subscale to measure the motivation deficits on behavioral activation) correlated positively with the horizontal axis (rho = 0.082, p = 0.009) but not the vertical (rho = 0.061, p = 0.055), while emotional apathy correlated positively with the vertical axis (rho = 0.097, p = 0.002) but not horizontal axis (rho = 0.040, p = 0.212). Moreover, GAD7 (i.e., generalized anxiety) correlated negatively with the horizontal manifold (rho = -0.071, p = 0.025) but not the vertical one (rho = -0.061, p = 0.054). No significant results were found for SHPS or PHQ-9 (all p > 0.06). All significant results reported here survive FDR correction for multiple comparisons (p < 0.05).

We considered whether position on the UMAP axes could be explained by combinations of symptoms, which might yield insight into their underlying meaning. We constructed a linear regression model to map several significant psychiatric symptoms on UMAP1 (horizontal) and UMAP2 (vertical), respectively. A linear regression model composed of UMAP1 as the dependent variable and anxiety, social apathy, and behavioral apath

y as independent variables showed a significant contribution of anxiety (t = -3.419, p < 0.001), as well as the two subscales from Apathy (social apathy, t = 2.854, p = 0.004; behavioral apathy, t = 2.156, p = 0.031). In contrast, a linear regression model with UMAP2 as the dependent variable and emotional apathy and social apathy as independent variables showed a significant contribution of emotional sensitivity (t = 2.529, p = 0.011) and a marginally significant contribution from social motivation (t = 1.784, p = 0.074) (Figure 4). We used variance inflation factor (VIF) to assess multicollinearity (21), and the results showed the VIF for both linear models is low (VIF < 5).

**Figure 4.**
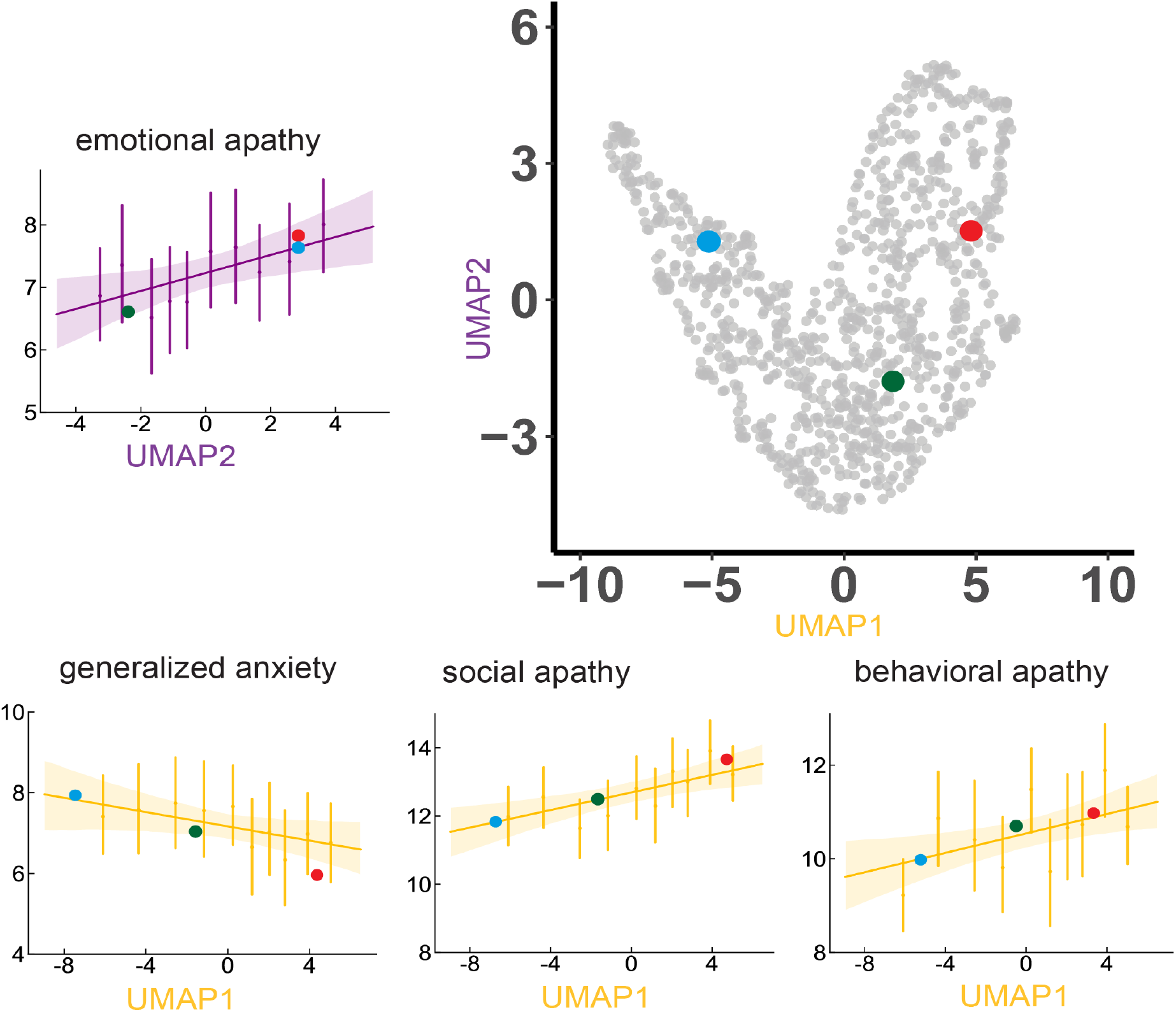
Manifold projection of individual differences in anxiety and apathy subdomains. Linear regression results demonstrate that levels of generalized anxiety, and social and behavioral apathy vary in opposite directions along the horizontal dimension UMAP1, while emotional apathy explains position along the vertical dimension UMAP2. The red, green, and blue dots are examples of different participants, showing how an individual’s position on the manifold is associated with their symptoms.

Traversing the manifold from the upper left, the propensity to explore gradually decreases, behavioral and social apathy increase, and anxiety decreases. At the same time the overall stability of states of both exploration and exploitation increases, and emotional apathy decreases. Moving through the trough of the manifold and up the right arm, while the probability of exploration continues to decrease, the decision states start to destabilize, and emotional apathy increases. Approaching the upper right, we find the highest level of exploitation, overall state instability, behavioral and social apathy, and the greatest emotional apathy.

## Discussion

Using dimensionality reduction on choice and reward sequences, we found that the distribution of individual differences in exploratory behavior in a 1001-person online experiment collapsed to a low-dimensional manifold. Computational modeling revealed that the stability and balance of decision-making states explained the axes of this manifold. Importantly, position on the manifold also predicted mental health symptoms that impact decision-making, revealing distinct roles for the transdiagnostic constructs of apathy and anxiety. These findings were consistent across linear and non-linear methods of dimensionality reduction (SI Section 5, Figure S2, Table S5&S6), robust to cross-validation (SI Section 6, Table S7&S8), and gender-independent (SI Section 7 & Figure S3).

Anxiety and apathy are often viewed as affective expressions of two opposing factors in the motivational organization of emotion: aversive and appetitive, respectively (22). While we might predict, then, that apathy and anxiety would be anti-correlated, in our large sample of 1001 people, they were instead positively correlated (r = 0.190, p < 0.001). However, when projected onto the behavioral manifold, we found that they did indeed oppose each other along the dimension controlling the exploration-exploitation balance. This counterintuitive result emerges from the fact that positive covariation of apathy and anxiety occurs independently of variation along the exploration-exploitation balance axis. The discovery of these independent axes of covariation resolves an apparent contradiction between the models of apathy and anxiety in the psychological literature, which align with prior work that anxious individuals are more exploratory (23), while apathetic individuals exhibit decision inertia and decreased exploration (24, 25) and the clinical neuropsychiatric literature (concordant with clinical experience) documenting their high degree of comorbidity (26, 27). The counterbalancing of apathy and anxiety we observe in exploration resembles the way that appetitive and aversive signals determine the balance between approach and avoidance. While approach and exploratory behavior may be responses to different kinds of environmental contexts and thus may provide differential advantages in those environments, they share common dopaminergic signaling pathways (28, 29), potentially explaining why they are related to similar affective states.

We found that the manifold axis corresponding to decision-state stability also explained individual differences in emotional apathy. Specifically, movement along this axis was associated with both a decrease in emotional *responsiveness* and a decrease in the persistence of decision-making states. One possible explanation is that the decreased reward sensitivity characteristic of emotional apathy also leads to a lowering of the activation energy barrier and the destabilization of decision-states. In line with this interpretation, a change in reward sensitivity is one of the most well-studied mechanisms by which emotions can affect decisions (30–32). While the decrease in both the win-stay and lose-shift behavior we observed along this same axis supports this conclusion, we did not observe a direct correlation between emotional apathy and exploratory behavior, suggesting that further work would be required to substantiate it.

Our findings bear on the ongoing quest to characterize psychiatric pathology through behavior, particularly when that pathology affects the kinds of flexible decision-making captured by the dynamics of exploration and exploitation. While recent work has challenged the notion that symptoms and behavior correspond in meaningful ways (33), we found that symptoms and behavior can be complementary in explaining the variation in biological markers of psychiatric state, with symptoms shaping how exploratory behavior relates to physiology (17). Consistent with this result, the present findings demonstrate that because variation in exploratory behavior follows a parabolically shaped manifold, co-variation in symptoms related to exploratory behavior, in turn, becomes shaped by this manifold, leading, for instance, to the opposing directions of apathy and anxiety, despite their positive correlation.

In summary, our results suggest that a generalizable latent structure links the balance and stability of exploration and exploitation decision strategies to apathy and anxiety. Although we examined normative behavior in an online population, one key implication of our findings is that in disorders characterized by cognitive rigidity, such as Parkinson’s Disease (34) or Schizophrenia (35), the relationship between behavior and affective symptoms may be similarly constrained. Locating an individual on the behavioral manifold might then help to predict how behavior would change with symptoms in response to treatment. For instance, the parabolic shape of the manifold with greater spread around its vertex suggests a potential “sweet spot” of balanced mild symptoms and flexible behavior that can be approached from different directions, depending on the starting point. To address this hypothesis, several important questions need to be answered: Do changes within an individual follow a trajectory on this same manifold? Do clinical populations project to the same manifold, or might they, in fact, be off manifold, projecting to the large regions of unoccupied space? The answers may help us to implement dimensional approaches to diagnosis and individualized neuropsychiatric care (36).

## Materials and Methods

### Ethics approval

The experimental procedures of all experiments were in line with the standards set by the Declaration of Helsinki and were approved by the local Research Ethics Committee of the University of Minnesota, Twin Cities. Participants provided written informed consent after the experimental procedure had been fully explained and were reminded of their right to withdraw at any time during the study.

### Participants

We recruited a sample of 1512 participants via Amazon Mechanical Turk (MTurk) as well as Prolific (Prolific.co), exclusion criteria included current or history of neurological, and psychiatric disorders. And 1001 participants completed all questionnaires and the bandit task (age: range 18-54, mean ± SD = 28.446 ± 10.354 years; gender, 493 female). All participants were compensated for their time in line with minimum wage.

### Questionnaire measurement

Participants’ depressive symptoms, anhedonia, general anxiety, and apathy symptoms were measured using the Patient Health Questionnaire (PHQ-9) (37), the Snaith-Hamilton Pleasure Scale (SHPS) (38), the General Anxiety Disorder Screener (GAD-7) (39), and the Apathy-Motivation Index (40), respectively. More specifically, the PHQ-9, a 9-item multiple-choice inventory, was employed to measure depressive symptoms. The SHPS, a 4-point Likert scale, measured symptoms of anhedonia. Participants’ general anxiety was measured using the GAD-7, which contains 7 items for assessing anxiety severity in the last two weeks. All items were rated on a 4-point scale, with higher scores indicating greater anxiety. Participants’ apathy level was measured using the 18-item Apathy-Motivation Index (AMI), which was designed to identify and measure the general apathy, as well as subtypes of apathy in behavioral, social, and emotional domains. Higher scores on AMI represent greater apathy. For all questionnaire scores, see SI, Section 1 & Table S1.

### Three-armed restless bandit task

We assessed exploration-exploitation behavioral dynamics using a 300-trial three-armed restless bandit task (18). Participants were free to choose between three targets for the potential to earn a reward of 1 point. Each target is associated with a hidden reward probability that randomly and independently changes throughout the task. We seeded each participant’s reward probability walks randomly to prevent biases due to particular kinds of environments. We assessed performance by comparing the total number of rewarded trials to that expected by chance. Out of the 1001 participants, 985 accrued more rewarded trials than would be expected by chance.

### Dimensionality reduction method

Popular and valid dimensionality reduction techniques to reveal manifolds include t-distributed stochastic neighborhood embedding (t-SNE) (41), uniform manifold approximation and projection (UMAP) (19), and Principal component analysis (PCA) (42). However, t-SNE suffers from limitations, including slow computation time, and loss of global data structure, and it is not a deterministic algorithm (43). The main drawback of PCA is that it is highly affected by outliers in the dataset (42). In contrast, UMAP is a deterministic and efficient algorithm, it also preserves both local and global structure of original high-dimensional data. Uniform Manifold Approximation and Projection (UMAP)

UMAP was implemented in the R language. The eight-dimensional datasets from all participants were passed into the R package *umap*, version 0.2.8.0 (19), available at https://cran.r-project.org/web/packages/umap/) with default parameter setting as n_component = 2, n_neighbors = 15, min_dist = 0.3, metric = ‘Euclidean’. For reproducibility reasons, we fixed the random_state in this algorithm. The hyperparameter *n_neighbors* decide the radius of the search region. Larger values will include more neighbors, thus forcing this algorithm to consider more global structure of original n-dimension data. Another important hyper-parameter, *min_dist* determines the allowed minimum distance apart for cases in lower-dimensional space. *metric* defines the way that UMAP is used to measure distances along the manifold.

### Model-free analyses

We adopted some widely used model-free measures, including win-stay and lose-shift (44, 45) as the direct measurement for this learning task.

*Win-stay*. Win-stay is defined as the percentage of times that the choice in trial *t*-1 was repeated on trial *t* following a reward.

*Lose-shift*. In contrast, lose-shift equals the percentage of trials that the choice was shifted or changed when the outcome of trial t-1 was non-reward.

### Computational Modeling

#### Hidden Markov Model

We fit a Hidden Markov Model (HMM) to the behavior, to decode the hidden state of each trial for each participant. Fundamentally, the HMM has two layers, the hidden layer (i.e., state) and observable layer. The hidden dimension should satisfy the Markov property. That is, the current hidden state only depends on the previous state but not any past model history. The observable dimension entirely depends on the current hidden states and is independent with other observations. Parameters of Hidden Markov Model can be represented as *Ω* → (*T, O, c*). Specifically, *T* is the transition probabilities matrix, *O* is the observation probabilities matrix, or emissions matrix, where the *c* refers to a vector with initial probabilities for each hidden state. Here, we have two hidden states, “exploration” state and the “exploitation” state.

The transition probability between exploration and exploitation states can be represented by:

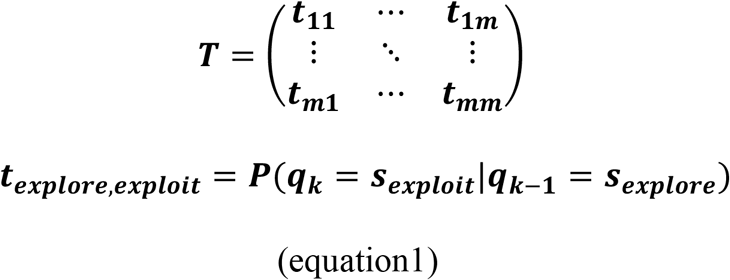

Where *t*_*explore,exploit*_ refers to the transition probability from hidden state *s*_*exploit*_ to another hidden state *s*_*explore*_

*k* = time instant, *m* = state sequence length

Then matrix *O* represent the transition probabilities between hidden and observable states.

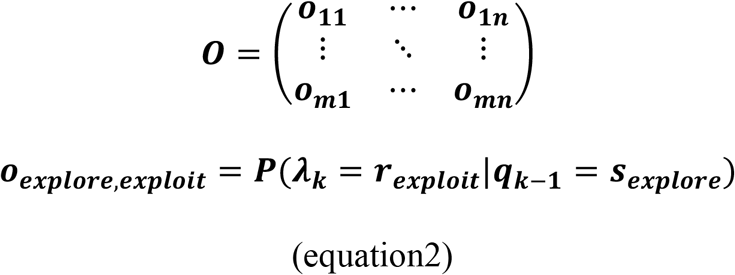

*k* = time instant, *n* = observation sequence length

Both matrix *O* and *T* satisfy the principle that the sum along the rows must be equal to 1.

*c* is a *m* dimensional row vector which refers to the initial probability distribution.

In our current study, the initial probability was fixed and equal to 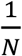, *N* equals the available choices.

We fit HMM via expectation-maximization using the Baum Welch algorithm and decode hidden state from observed choice sequences by the Viterbi algorithm (46).

#### Analyzing model dynamics

When we successfully decode the state sequence from HMM, the next question would be, what is the long-term probability that the system will be in either state of exploration or exploitation? To answer this question, we further extracted the steady state analysis from the hidden Markov process from the transition matrix. Let π⇒ steady-state probabilities, mathematically,

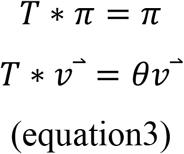

Where 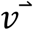 is the eigenvector and *θ* is the eigenvalue, *T* is the transition matrix again. The steady-state vector is the eigenvector with eigenvalue 1.

To develop an intuition for how psychiatric symptoms covary with the dynamics of exploration-exploitation decision-making, we describe the energetic landscape of behavior (20). Each state has an energy associated with it. Like a valley between two high peaks, a low-energy state is very stable and deep, while a high-energy condition is less stable. The steady-state probability is related to the energy of that state via Boltzmann distribution:

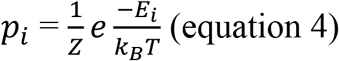

where *Z* is the partition function of the system, *k*_*B*_ is the Boltzmann constant, and *T* is the temperature of the system (a constant). To characterize the difference in energy associated with *exploration* and *exploitation*, we have:

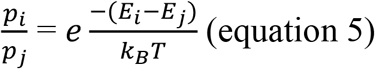

after rearranging, we will have:

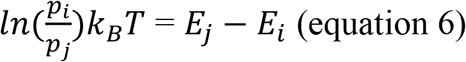

Next, we are going to quantify the activation energy required to transition between exploration and exploitation. We adopted the Arrhenius equation from chemical kinetics that relates the transitional probabilities between different states to the energy required to alert these transitions. The activation energy required to escape state (*E*_*a*_):

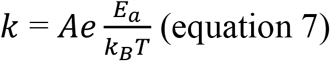

where the *k* is the transition probability from one state to another state (e.g., from exploitation to exploration), *E*_*a*_ represents the activation energy required to escape that state (e.g., exploitation) where *A* is a constant and we set this to one for convenience.

after rearranging, we will have:

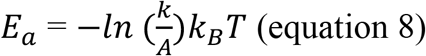

## Acknowledgements

This work was supported by NIMH under award #R21MH127607, NIDA under award #K23DA050909, and the University of Minnesota’s MnDRIVE (Minnesota’s Discovery, Research and Innovation Economy) initiative.

We thank A. David Redish, Cathy Chen, Seth D Koenig, and Brian Sweiss for helpful comments on the manuscript.

## Code and data availability

The data, and source code used to do data analysis, as well as generate main and supplementary figures, is published at https://github.com/hermandarrowlab/umap-bandit

## Notes

### Competing Interest Statement

The authors have declared no competing interest.

### Summary of Updates

Fixed some errors, slight rewording for clarity

