## Supplemental Material for "A low dimensional manifold of human exploratory behavior reveals opposing roles for apathy and anxiety"

### A low dimensional manifold of human explore/exploit behavior shapes states of apathy and anxiety

* Corresponding author: Alexander B. Herman

**This PDF file includes:**

Supporting text

8 SI Sections

Figures S1 to S4

Tables S1 to S8

### Supplemental information

**Section1. Descriptive statistic for all questionnaires in current study**

We reported the basic statistical information for all questionnaires in current study in Table S1.

**Table S1. Descriptive statistics for all questionnaires.**

| | PHQ-9 | GAD-7 | SHPS | Apathy | Apathy-ES | Apathy-BA | Apathy-SM |
|---|---|---|---|---|---|---|---|
| Mean | 9.53 | 7.17 | 1.92 | 30.46 | 10.54 | 12.69 | 7.23 |
| SD | 6.13 | 5.54 | 2.26 | 9.32 | 5.04 | 4.80 | 4.22 |
| Minimum | 0 | 0 | 0 | 4 | 0 | 0 | 0 |
| Maximum | 28 | 21 | 14 | 64 | 24 | 24 | 24 |

[a]PHQ-9 = Patient Health Questionnaire; SHPS = Snaith-Hamilton Pleasure Scale; GAD-7 = General Anxiety Disorder Screener; Apathy-BA = Apathy behavioral activation; Apathy-ES = Apathy emotional sensitivity; Apathy-SM = Apathy social motivation.

#### Section2. Manifold from 32-dimension dataset

We next considered whether including additional reward history in our analysis would affect the results from dimensionality reduction. We found that the resultant manifold shape is similar to the manifold revealed in the eight-dimension dataset. We extended the encoding of trial-by-trial behaviors to two trials before the current $t^{th}$ trial (three consecutive trials) {$choice_{t-2}$, $outcome_{t-2}$, $choice_{t-1}$, $outcome_{t-1}$, $choice_t$}, resulting in a dataset with thirty-two dimensions. However, we found a sparse distribution of data in these thirty-two dimensions, with many zero-valued behavioral bins, suggesting the thirty-two-dimension dataset contains decision patterns that happen only rarely (e.g., stay ~ non-reward ~ stay ~ non-reward ~ stay; switch ~ non-reward ~ stay ~ reward ~ switch). Nonetheless, we found that the resultant manifold shape is similar to the manifold revealed in the eight-dimension dataset. As a result, we proceeded with further analysis of the two consecutive trial eight-dimensional data.

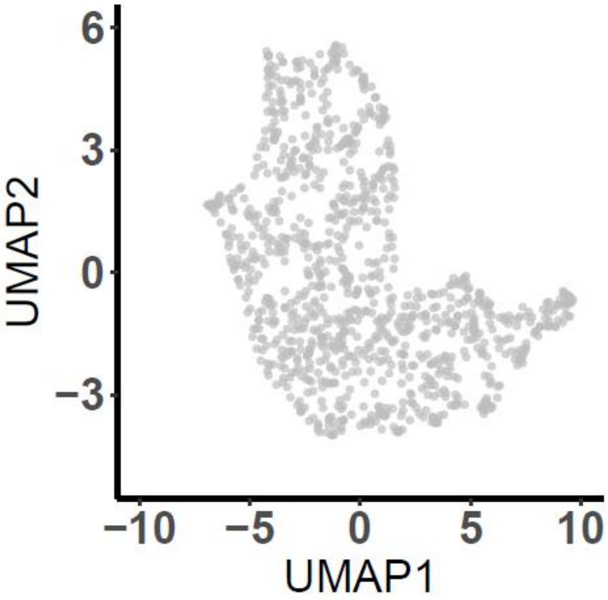


**Figure S1. Manifold from a 32-dimension dataset.**

#### Section 3. Model-free and model-based results

**Table S2A**

| | Model-free indices | | | |
|---|---|---|---|---|
| | Stay % | Switch % | Win. Stay | Lose.shift |
| Mean | 0.646 | 0.353 | 0.861 | 0.732 |
| SD | 0.180 | 0.180 | 0.210 | 0.204 |

**Table S2B**

| | Model-based indices | | | | | |
|---|---|---|---|---|---|---|
| | Exploitation% | Exploration% | Exploit-Exploit | Exploit-Explore | Explore-Explore | Explore-Exploit |
| Mean | 0.544 | 0.455 | 0.811 | 0.188 | 0.818 | 0.181 |
| SD | 0.243 | 0.243 | 0.215 | 0.215 | 0.146 | 0.146 |

**Section4. Correlation between model-free indices and UMAP score, as well as model-based indices.**

**Table S3**

| | GAD7 | Apathy -total | Apathy -BA | Apathy -SM | Apathy -ES | UMAP1 | UMAP2 |
|---|---|---|---|---|---|---|---|
| Switch% | 0.077* | -0.123*** | -0.084** | -0.118*** | -0.037 | -0.920*** | -0.107*** |
| Stay% | -0.077* | 0.123*** | 0.084** | 0.118 | 0.037 | 0.920*** | 0.107*** |
| Win.stay | -0.037 | 0.072* | 0.053 | 0.091** | -0.007 | 0.801*** | -0.281*** |

| | | | | | | | |
|---|---|---|---|---|---|---|---|
| Lose.shift | 0.109*** | -0.146*** | -0.092** | -0.103** | -0.096** | -0.458*** | -0.746*** |

[a] *** p < 0.001; ** p < 0.01; * p < 0.05. All significant P-value reported here can be survived after FDR correction (original Benjamini & Hochberg FDR procedure, threshold, p<0.05). Apathy-BA = Apathy behavioral activation; Apathy-ES = Apathy emotional sensitivity; Apathy-SM = Apathy social motivation.

**Table S4**

| | GAD7 | Apathy -total | Apathy -BA | Apathy -SM | Apathy -ES | UMAP1 | UMAP2 |
|---|---|---|---|---|---|---|---|
| Exploration % | 0.061 | -0.128*** | -0.093** | -0.122*** | -0.033 | -0.902*** | -0.194*** |
| Exploitation % | -0.061 | 0.128*** | 0.093** | 0.122*** | 0.033 | 0.902*** | 0.194*** |
| Exploit-exploit | -0.047 | 0.044 | 0.047 | 0.053 | -0.019 | 0.605*** | -0.222*** |

| | | | | | | | |
|---|---|---|---|---|---|---|---|
| Explore-explore | 0.021 | -0.118*** | -0.067[#] | -0.133*** | -0.029 | -0.418*** | -0.391*** |
| Explore-exploit | -0.021 | 0.118*** | 0.067[#] | 0.133*** | 0.029 | 0.418*** | 0.391*** |
| Exploit-explore | 0.047 | -0.044 | -0.047 | -0.053 | 0.019 | -0.605*** | 0.222*** |

[a] *** $p < 0.001$; ** $p < 0.01$; * $p < 0.05$. All significant P-value reported here can be survived after FDR correction (original Benjamini & Hochberg FDR procedure, threshold, $p<0.05$). Apathy-BA = Apathy behavioral activation; Apathy-ES = Apathy emotional sensitivity; Apathy-SM = Apathy social motivation.

[#] these p-values cannot survive after FDR-correction ($p<0.05$)

#### Section 5. Manifold from t-SNE and PCA showed similar results

We conducted t-Distributed Stochastic Neighbor Embedding (t-SNE) and Principal Component Analysis (PCA) to confirm the manifold. The same eight-dimensional datasets from all participants were passed into the R package Rtsne, version 0.16 (available at https://cran.r-project.org/web/packages/Rtsne/index.html) with default parameter setting as n_component = 2, perplexity = 30, min_iter = 1000, metric = 'Euclidean'. We showed a similar manifold shape as UMAP found. Like in the main text, we also mapped model-free indices, as well as parameters from HMM onto t-SNE manifolds. The meaning of gradient change here is the same as with the UMAP manifold. We further did a Spearman correlation between t-SNE score and psychiatric symptoms, the results were quite similar. That is, t-SNE1 was positively correlated with Apathy (rho = 0.133, $p < 0.001$), as well as its subscale social apathy (rho = 0.123, $p < 0.001$), behavioral apathy (rho = 0.093, $p = 0.003$), but not emotional apathy (rho = 0.053, $p = 0.089$), and t-SNE1 negatively correlated with GAD7 (rho = -0.078, $p = 0.013$). t-SNE2, however, negatively correlated with apathy on emotional apathy (rho = -0.086, $p = 0.006$) but not correlated with another two dimensions of apathy (all $p > 0.330$).

Robustly, the low-dimensional space from PCA is also quite similar to the manifold from UMAP and t-SNE. Analyzing Spearman's correlations between PCA score and psychiatric symptoms also leads to the same conclusion. That is, the horizontal axis of low-dimensional space positively correlated with apathy (rho = 0.138, $p < 0.001$) but negatively correlated with anxiety (rho = -0.081, $p = 0.010$). Again, the horizontal axis positively correlated with subscales of apathy including social apathy (rho = 0.131, $p < 0.001$), behavioral apathy (rho = 0.096, $p = 0.002$), but not emotional apathy (rho = 0.058, $p = 0.065$), while the vertical one only correlated with emotional apathy (rho = -0.101, $p = 0.001$)

All other correlation results based on t-SNE and PCA scores can be found in tables below.

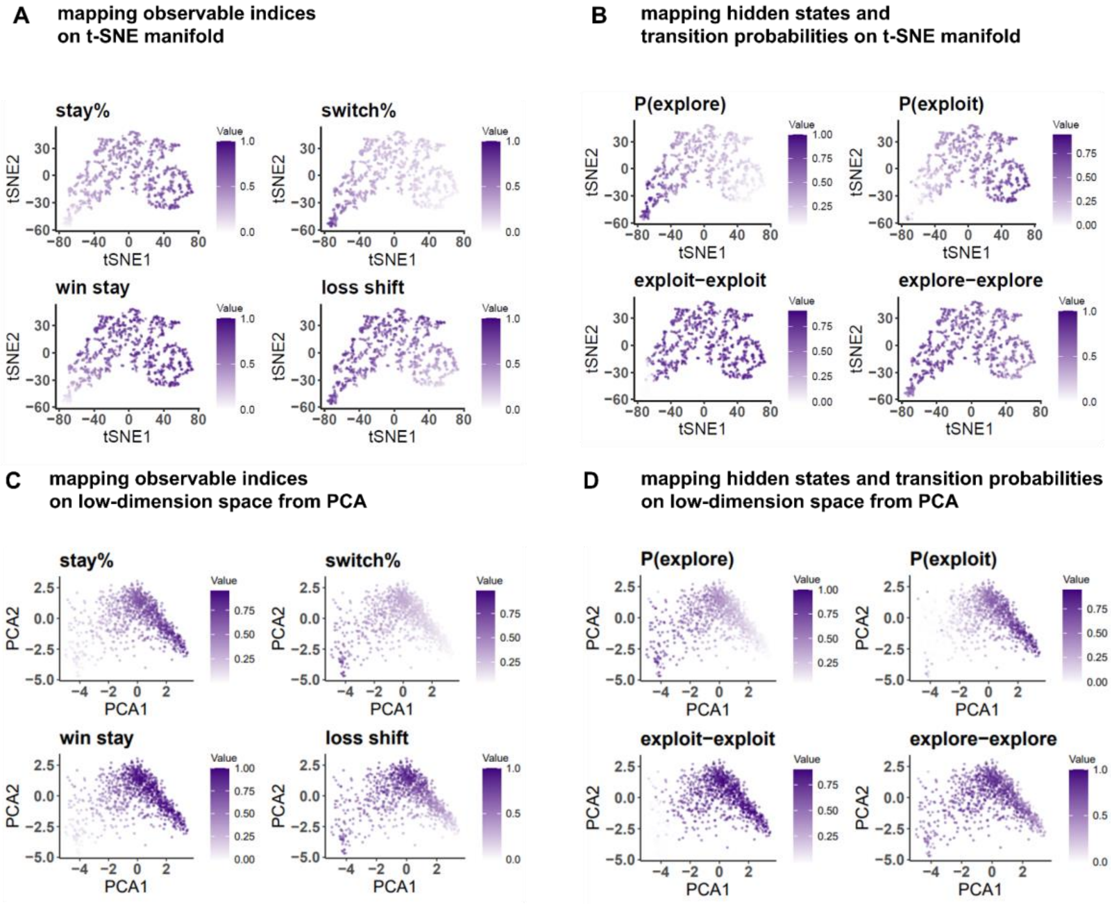


**Figure S2. Manifold from t-SNE and PCA and its implications for behavioral tasks.**

Both gradient maps are similar to the manifold from UMAP. Statistical results see Table S5 (for model-free indices) and Table S6 (for model-based parameters of Hidden Markov Model)

**Table S5**

| | t-SNE1 | t-SNE2 | PCA1 | PCA2 |
|---|---|---|---|---|
| Switch% | -0.914*** | -0.323*** | -0.967*** | -0.139*** |
| Stay% | 0.914*** | 0.323*** | 0.967*** | 0.139*** |
| Win.stay | 0.713*** | 0.608*** | 0.832*** | 0.513*** |
| Lose.shift | -0.631*** | 0.482*** | -0.665*** | 0.603*** |

[a] *** $p < 0.001$; ** $p < 0.01$; * $p < 0.05$. All significant P-values reported here survive FDR correction.

(Original Benjamini & Hochberg FDR procedure, $p < 0.05$)

**Table S6**

| | t-SNE1 | t-SNE2 | PCA1 | PCA2 |
|---|---|---|---|---|
| Exploration % | -0.912*** | -0.250*** | -0.937*** | -0.016*** |
| Exploitation % | 0.912*** | 0.250*** | 0.937*** | 0.016 |
| Exploit-exploit | 0.536*** | 0.434*** | 0.683*** | 0.357*** |
| Explore-explore | -0.484*** | 0.111*** | -0.449*** | 0.199*** |
| Explore-exploit | 0.484*** | -0.111*** | 0.449*** | -0.199*** |
| Exploit-explore | -0.536*** | -0.434*** | -0.683*** | -0.357*** |

[a]all significant P-values reported here survive FDR correction.

(Original Benjamini & Hochberg FDR procedure, $p<0.05$)

#### Section6. Results from validation analysis

To prove the manifold is valid and robust, we conducted a 10-fold cross-validation analysis by creating 10 randomly trained samples (N=667), each consisting of two-thirds of the full dataset, and corresponding 10 randomly tested samples (N=334). Specifically, we projected the test sample onto the UMAP embedding based on the training sample and then compared it (Pearson correlation analysis) with the UMAP manifold generated from the test sample itself. We repeated this procedure 10 times and then averaged the correlation coefficients to measure the accuracy of the prediction. Results showed that the prediction accuracy for UMAP1 is 94.37% while the prediction accuracy for UMAP2 is 76.99%. These high correlation coefficients confirmed the robustness of the manifolds we have.

We further constructed the same linear regression analysis as did on the full dataset. For results, see TableS7 and TableS8. Although the correlation direction between psychiatric symptoms and UMAP properties is different among resampled datasets, the conclusions are the same. That is, individuals who lack more motivation on social and behavioral apathy would show more stay behaviors and more exploitation states, while individuals who showed higher anxiety levels would exhibit more switch behaviors and exploration states.

**Table S7. Linear regression results based on a resampled dataset for UMAP1.**

| | 1 | 2 | 3 | 4 | 5 | 6 | 7 | 8 | 9 | 10 |
|---|---|---|---|---|---|---|---|---|---|---|
| (Intercept) | 0.41 | 0.98 ** | -0.78 ** | -0.60 * | 0.73 ** | -0.70 ** | -1.07 *** | 0.17 | -0.65 * | 0.50 |
| | (0.26) | (0.31) | (0.30) | (0.24) | (0.28) | (0.27) | (0.29) | (0.21) | (0.27) | (0.29) |
| ApathyES | -0.02 | -0.03 | 0.05 * | 0.02 | -0.05 * | 0.03 | 0.05 * | -0.02 | 0.02 | -0.01 |
| | (0.02) | (0.02) | (0.02) | (0.02) | (0.02) | (0.02) | (0.02) | (0.02) | (0.02) | (0.02) |
| ApathySM | -0.03 * | -0.07 *** | 0.04 * | 0.04 ** | -0.03 | 0.04 * | 0.06 *** | -0.02 | 0.05 ** | -0.04 * |
| | (0.02) | (0.02) | (0.02) | (0.02) | (0.02) | (0.02) | (0.02) | (0.01) | (0.02) | (0.02) |

[a] *** $p < 0.001$; ** $p < 0.01$; * $p < 0.05$.

The number in the first row represents the number of the resampled dataset (e.g., 1 = resample 1). Coefficients and standard error showed (stand error showed in parentheses). VIF for all models is smaller than 5.

**Table S8. Linear regression results based on resampled dataset for UMAP2.**

| | 1 | 2 | 3 | 4 | 5 | 6 | 7 | 8 | 9 | 10 |
|---|---|---|---|---|---|---|---|---|---|---|
| (Intercept) | -1.19 ** | 0.77 | 0.55 | -0.97 * | 1.23 ** | 1.33 ** | -0.90 * | 1.03 | 1.27 ** | 0.85 * |
| | (0.45) | (0.39) | (0.38) | (0.46) | (0.41) | (0.43) | (0.42) | (0.53) | (0.47) | (0.39) |
| GAD7 | -0.09 ** | 0.06 * | 0.05 * | -0.08 ** | 0.06 * | 0.06 * | -0.10 *** | 0.11 ** | 0.07 * | 0.03 |
| | (0.03) | (0.02) | (0.02) | (0.03) | (0.03) | (0.03) | (0.03) | (0.03) | (0.03) | (0.03) |
| ApathySM | 0.11 ** | -0.04 | -0.03 | 0.06 | -0.10 *** | -0.08 * | 0.08 * | -0.09 * | -0.07 * | -0.05 |
| | (0.04) | (0.03) | (0.03) | (0.04) | (0.03) | (0.03) | (0.03) | (0.04) | (0.04) | (0.03) |
| ApathyBA | 0.05 | -0.06 | -0.05 | 0.08 * | -0.03 | -0.07 * | 0.06 | -0.06 | -0.08 * | -0.04 |
| | (0.04) | (0.03) | (0.03) | (0.03) | (0.03) | (0.03) | (0.03) | (0.04) | (0.03) | (0.03) |

[a] *** $p < 0.001$; ** $p < 0.01$; * $p < 0.05$. The number in the first row represents the number of the resampled dataset (e.g., 1 = resample 1). Coefficients and standard error showed (stand error showed in parentheses). VIF for all models is smaller than 5.

#### Section 7. Gender difference does not affect manifolds' psychiatric meaning

To test whether there is a gender effect on manifold representation and its association with psychiatric symptoms, we compared the two-dimensional UMAP score between females and males, as well as mapping the gender label onto the UMAP manifold (Figure S3), then did the same regression analysis for each gender group and test the difference statistically.

According to Figure S3, the gender label did not show an obvious difference on the UMAP1, which indicates that there is no gender difference in manifold representation. This conclusion can be further assisted by an independent T-test on the UMAP1 (UMAP1 score, $t(987)^{a} = -0.267$, $p = 0.789$). But statistically, we found significant gender differences on UMAP2 (UMAP2 score, $t(987) = -3.594$, $p<0.001$). Next, we constructed a linear regression model with gender effect (lm package in R). Unsurprisingly, all interactions with gender were not significant (all $p > 0.300$).

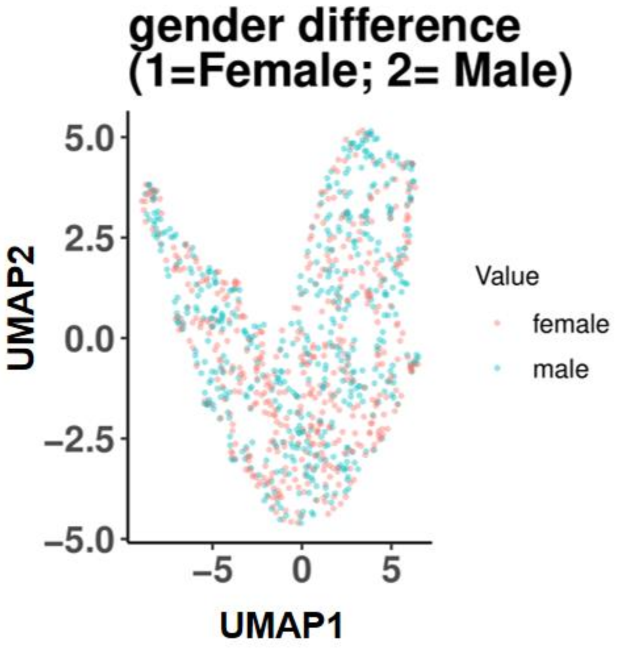


**Figure S3. Mapping gender labels onto the manifold.**

[a] We have 12 participants who reported their gender as "other", so we excluded these 12 participants when we discussed gender related results.

**Section 8. Manifold from random choice**

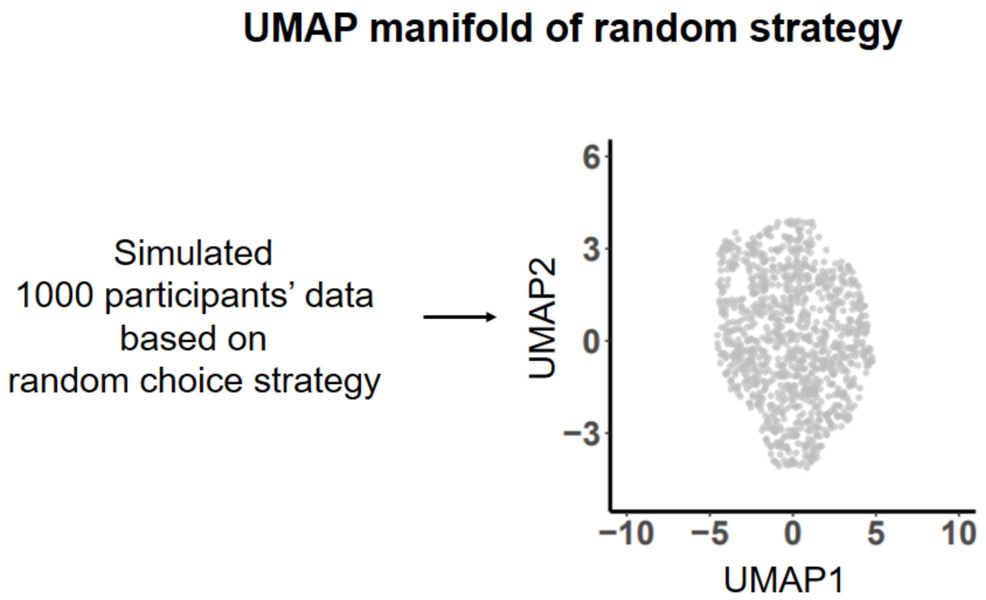


**Figure S4. Manifold derived from simulated data, based on random choice.**
